# Solving musculoskeletal biomechanics with machine learning

**DOI:** 10.1101/2020.08.24.263962

**Authors:** Yaroslav Smirnov, Denis Smirnov, Anton Popov, Sergiy Yakovenko

**Affiliations:** Department of Electronic Engineering, Igor Sikorsky Kyiv Polytechnic Institute, Kyiv, Ukraine; Department of Computer-aided Management and Data Processing Systems, Igor Sikorsky Kyiv Polytechnic Institute, Kyiv, Ukraine; Data & Analytics, Ciklum, London, UK; Department of Human Performance, School of Medicine, West Virginia University, Morgantown, WV, USA

## Abstract

Deep learning is a relatively new computational technique for the description of the musculoskeletal dynamics. The experimental relationships of muscle geometry in different postures are the high-dimensional spatial transformations that can be approximated by relatively simple functions, which opens the opportunity for machine learning applications. In this study, we challenged general machine learning algorithms with the problem of approximating the posture-dependent moment arm and muscle length relationships of the human arm and hand muscles. We used two types of algorithms, light gradient boosting machine (LGB) and fully connected artificial neural network (ANN) solving the wrapping kinematics of 33 muscles spanning up to six degrees of freedom (DOF) each for the arm and hand model with 18 DOFs. The input-output training and testing datasets were generated by our previous phenomenological model based on the autogenerated polynomial structures (Sobinov et al., 2019). Both models achieved a similar level of errors: ANN model errors were 0.08±0.05% for muscle lengths and 0.53±0.29% for moment arms, and LGB model made similar errors—0.18±0.06% and 0.13±0.07%, respectively. LGB model reached the training goal with only 10^3 samples, while ANN required 10^6 samples; however, LGB models were about 39 slower than ANN models in the evaluation. The sufficient performance of developed models demonstrates the future applicability of machine learning for musculoskeletal transformations in a variety of applications, such as in advanced powered prosthetics.

**Author Summary:** The accurate decoding of arm and hand motor intent from biological signals remains a key challenge. Solving this task with machine learning requires vast posture- and task-dependent data for identifying structural and functional parameters within dynamic musculoskeletal relationships. This problem is related to *the curse of dimensionality* where the processing complexity grows exponentially with the number of degrees of freedom described by the model. Here, we developed a tool based on artificial neural networks (ANN) to solve the kinematic transformation from posture to muscle path length and muscle moment arms. We used an accurate model of posture-dependent muscle moment arms and length to train and test the ability of ANN to solve this high-dimensional and computationally intense transformation and compare it to the boosted decision tree approach. We demonstrated that model-driven training is an efficient method to handle the encoding of high-dimensional musculoskeletal relationships. Adding muscles to the transformation, which increases the input-output complexity, does not reduce the prediction accuracy and does not require the increase in the number of elements within the network demonstrating the viability of this approach for applications using musculoskeletal biomechanics.

## Introduction

Machine learning (ML) with artificial neural networks (ANN) is revolutionizing applications where recognition, cognition, and categorization abilities used to require direct human involvement (LeCun et al., 2015). While the initial focus was on the substitution of human operators, for example, in driving a vehicle (Shi et al., 2020), classifying media (Gupta and Katarya, 2020), or for radiological diagnostics of benign or malignant mass in mammography (McKinney et al., 2020; Shen et al., 2019), the reach of ANN extended to the exploration of sensorimotor mechanisms (Richards et al., 2019). The structural and functional complexity of this system, which is distributed across multiple neural and mechanical pathways (Valero-Cuevas, 2015) and has high-dimensional computations for our segmented body control (Bernstein, 1967), offers a unique challenge and opportunity for this approach. The opportunity lies within the embedded ability of ANNs to absorb and classify a high volume of multidimensional input-output data signals, which are symptomatic of motor processing responsible for coordinated spatiotemporal action of multiple muscles generating movement. Yet another extreme expression of the neural processing complexity and efficiency is the brain’s ability to solve the Bernsteinian degrees of freedom problem where the same motor goal of body control can be accomplished with different kinematic solutions (Bernstein, 1967). ML methods based on ANNs can potentially resolve or, at least, identify targets for the long-standing theoretical challenge that can provide insight to the current theories of neural processing (McNamee and Wolpert, 2019) and has multiple practical human-machine applications, *e*.*g*., in advanced prosthetics (Dantas et al., 2019).

Developing a fast and intuitive interface with a high-dimensional artificial limb is a challenging task solved increasingly with the help of pattern recognition algorithms (Geethanjali, 2016a; Muceli and Farina, 2012). Typically, myoelectric direct and proportional control is used to decode motor intent from the recorded surface electromyography (EMG and convert it into torques or positions of the powered prosthetic devices (Geethanjali, 2016b). This was previously accomplished using ANN decoding algorithms decoding kinematics from EMG signals (Muceli and Farina, 2012). The intuitive control requires an additional transformation based on the representation of the controlled device and its neural control (Castellini and van der Smagt, 2009). The failure to account for the dynamics of prosthesis would lead to direct kinematic errors. The failure to recognize the biological strategies in solving limb dynamics would reduce robustness and intuitiveness of control even when the control of mechanical devices is perfectly tuned. The latter would occur because the interlimb inertial dynamics is encoded within neural commands even when limb dynamics changes. For example, mechanical shoulder immobilization does not abolish the stabilizing shoulder muscle activity during elbow movement (Debicki and Gribble, 2005; Gribble and Ostry, 1999, 1998). The expected musculoskeletal dynamics persists within neural commands months and years after the acute stage of limb trauma and amputations. The successful use of these commands for prosthetic control would theoretically require the representation of musculoskeletal and segmental limb dynamics to account for the dynamics encoded within neural control signals.

Relatively few studies applied ML techniques to musculoskeletal dynamics. The early applications of simple feed-forward ANNs allowed mapping of an average locomotor pattern of 16 EMGs to hip, knee, and ankle joint angles and moments (Sepulveda et al., 1993). Even though the muscle paths and muscle force generation were not simulated in this study, the decoding was demonstrated with low errors. The changes in the locomotor pattern at slow and fast speeds can also be generalized by a simple ANN with supervised learning via backpropagation (Heller et al., 1993). Furthermore, the mapping can be done not only between the locomotor activation patterns, but also with the muscle forces; albeit, the accurate predictions within trials have not been demonstrated (Liu et al., 1999). Yet, similar type of statistical mapping, admittedly, can be expressed with standard dimensionality reduction techniques with high precision and low computational cost, *e*.*g*., principal component analysis (PCA) (Patla, 1985). Multiple methodological variations have been since developed and applied to solve musculoskeletal problems. Notably recurrent ANNs were used to predict elbow torques from EMGs and showed the benefit of taking into account kinematic inputs, joint angle and velocity (Song and Tong, 2005). A combination of convolutional and recurrent ANNs can accurately and robustly map from the time-frequency frames of multi-channel EMG to limb movement (Xia et al., 2018). Purely statistical learning of the musculoskeletal transformation from posture to the control inputs has been also demonstrated with multiple hybrid ANN methods for a 7 DOF robotic arm with artificial muscles (Hartmann et al., 2013). Similar to our approach the musculoskeletal transformation was learnt from an input-output dataset. While the tracking of two DOF simple arm was achieved, the control of both robot and its simulation resulted in large tracking errors. The high-dimensional control remains a challenge.

While data-driven mapping with ANNs or using PCA and other statistical classification alternatives are robust, these methods do not generally capture the mechanistic relationships. Intrinsic muscle properties and musculoskeletal organization contribute to the muscle force generation, and these mechanistic details may assist in the reconstruction of relationships between neural commands and generated movement. In this study, we aim to challenge the problem of learning the musculoskeletal dynamics (MSD) with several simpler machine learning techniques, looking for a more computationally efficient solution. MSD requires high-dimensional transformations of posture into muscle moment arms and length, which are the essential variables in the calculation of generated muscle forces. The Hill-type muscle model can then allow us to define the posture-dependent force-length-velocity dependency (Figure 1, Muscle Model) (Zajac, 1989) and next to compute muscle and joint torques. The remaining step for the generation of movement is the simulation of equations of motion by using a physics engine. The critical constraint for the accuracy of these computations is the loop latency that limits the computational stability of integration (Todorov et al., 2012). Thus, the goal for real-time biomechanics is to implement a method with high accuracy and low computational cost of musculoskeletal transformations. In this study, we solve the problem of mapping the joint posture expressed with angles to muscle moment arms and their muscle lengths. We developed the training and validation of two machine learning methods based on the model-driven generation of the input-output dataset for arm and hand kinematics. We present the validation and performance metrics focused on the goal of developing the musculoskeletal transformation for online control problems.

**Figure 1.**
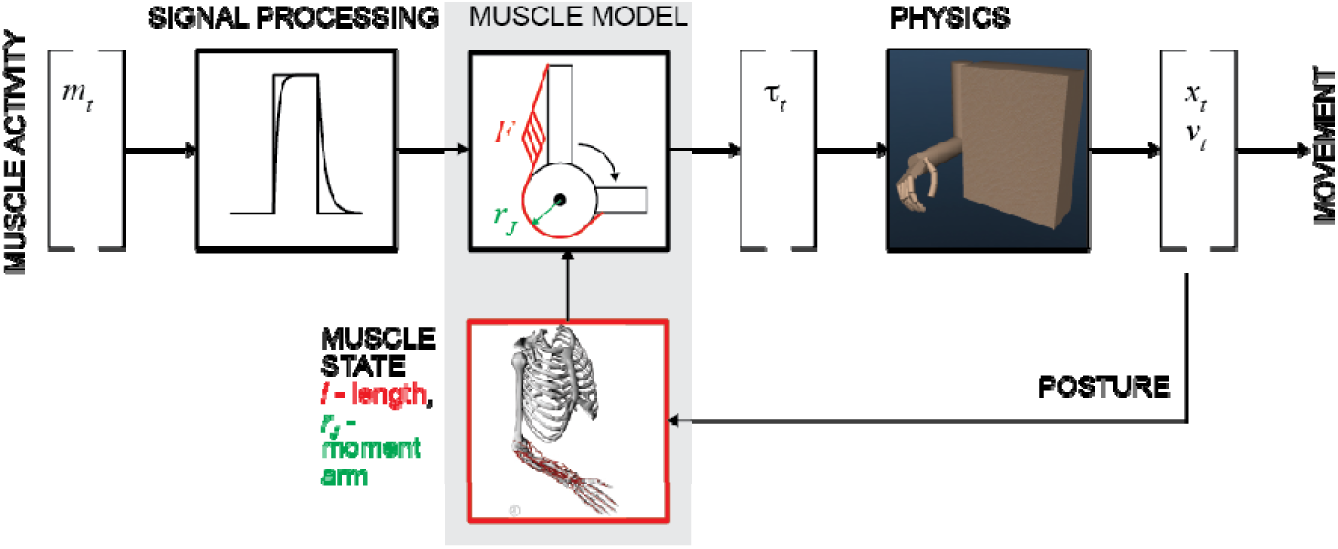
General concept of motor intent decoding from muscle activity. The schematic illustrates the transformation of EMG inputs through signal processing and musculoskeletal relationships (muscle model) into estimated torques that actuate limbs to generate movement by solving the equations of motion (physics). Limb posture modifies nonlinear muscle force-length-velocity relationship and torques.

## Methods

### Musculoskeletal Polynomial Model for Generation of Training and Testing Datasets

We have previously developed the method of autogenerated polynomial models (Sobinov et al., 2019). In these polynomials, the composition of terms was expanded using objective information measurements, i.e., the corrected Akaike Information Criterion. In brief, the posture-dependent musculotendon actuator length and joint moment arms for each muscle in the upper-limb model (Gritsenko et al., 2016) were accurately approximated using the selection of up to 5th power polynomial terms, where muscle length and moment arms were connected through a partial derivative of the muscle length in local coordinates corresponding to limb posture. Overall, the 18 DOF model of the human arm and hand is actuated by 33 muscles, each spanning about 3 DOFs and up to 6 DOFs for thumb muscles. Thus, each actuator is represented by a set of one length- and about 3 moment arm-posture polynomials. The goal of this development was to bypass costly calculations of geometrical transformations with high-quality approximations. Previously, we have demonstrated high-fidelity of these approximations with kinematic errors below 1% (Sobinov et al., 2019). These polynomial models of muscle posture-dependent state were used to develop an ANN-based approximation method for the musculoskeletal dynamics in this study.

### Training, Validation, and Testing Datasets

Training, validation, and testing datasets for the assessment of model performance were generated by the musculoskeletal polynomial model, which was used as a reference in this study. Input-output relationships were extracted randomly with uniform distribution where inputs were 18 DOF vectors of joint angles and the outputs were 33 length vectors and 99 moment arm vectors. An average muscle crosses 3 DOFs and has, consequently, 3 moment arm relationships on average. We used the supervised learning approach for training ML models. The training dataset was used for two tasks, tuning the model hyper-parameters and model training, to maximize the model performance in replicating the desired outputs with given inputs. The testing dataset contained ∼5% of all data (5*10^4 samples). The remaining ∼95% were divided into the training dataset (80%, 8*10^5 samples) and the validation dataset (20%, 2*10^5 samples). The validation dataset was used to prevent overfitting, i.e. higher performance on the training data as compared to that on the validation data. These datasets were similarly used for training ANN and LGB models, described below.

### Metrics

The performance of the trained models was further evaluated with the testing dataset, which was not used during the training procedure. We expected to reach the same error tolerances as in our previous polynomial fitting method study (Sobinov et al., 2019). Consequently, we used the same normalization of lengths and moment arms as in our previous work. The RMSE values were calculated as the absolute difference between reference and predicted muscle length values. To normalize the results we divided each reference and predicted length value by the muscle length range respectively:

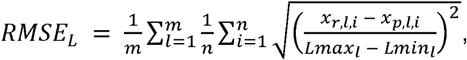

where *m* was the number of muscles (*m*=33), *n* was the number of test samples, *x*_*r,l,i*_ and *x*_*p,l,i*_ were reference and predicted length values, respectively, *Lmax*_*l*_ and *Lmin*_*l*_ were the maximum and minimum values over the full range of *l*th muscle length.

Similarly, the RMSE of moment arms was calculated as the absolute difference between reference and predicted values, which were normalized to the moment arm maximum (*Mmax*_*l*_):

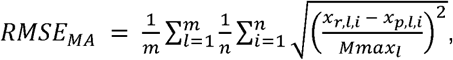

where *x*_*r,l,i*_ and *x*_*p,l,i*_ were reference and predicted values, *m* was the number of moment arms (*m*=99), and *n* was the number of test samples.

### Machine Learning Models

Two types of ML models were used to map the musculoskeletal input-output relationships. We used the light gradient boosting machine (LGB) models and artificial neural network (ANN) with two hidden layers. The models were trained and tested according to the workflow in Fig.2A. Validation accompanied the training process to prevent overfitting.

#### Light gradient boosting machine (LGB)

LGB algorithms belong to the group of gradient boosting methods based on choosing iteratively simple learner functions that point to the global minimum in the cost function. Gradient boosting is a technique to assemble weak prediction models, in our case regression trees, as processing stages that reduce performance errors. Here, the regression trees use binary recursive decisions to follow a path alone hierarchically organized nodes that terminate with the final branches, called leaves. The training process was the search for the optimal routing of inputs so that similar outputs were grouped together (Dantas et al., 2019; Microsoft Corporation, n.d.). The boosting method assembles the sequences of multiple regression trees to process errors in stages and gradually improve output accuracy (Fig. 2B). We used gradient-based one-side sampling in LGB to select a set of inputs where previous weak learner models have the largest output errors. The structure of the decision trees adapted to the required error tolerance by expanding the number of nodes (leaves) up to the maximal preset value determined empirically. We used Microsoft open source implementation of LGB (Microsoft Corporation, n.d.).

**Figure 2.**
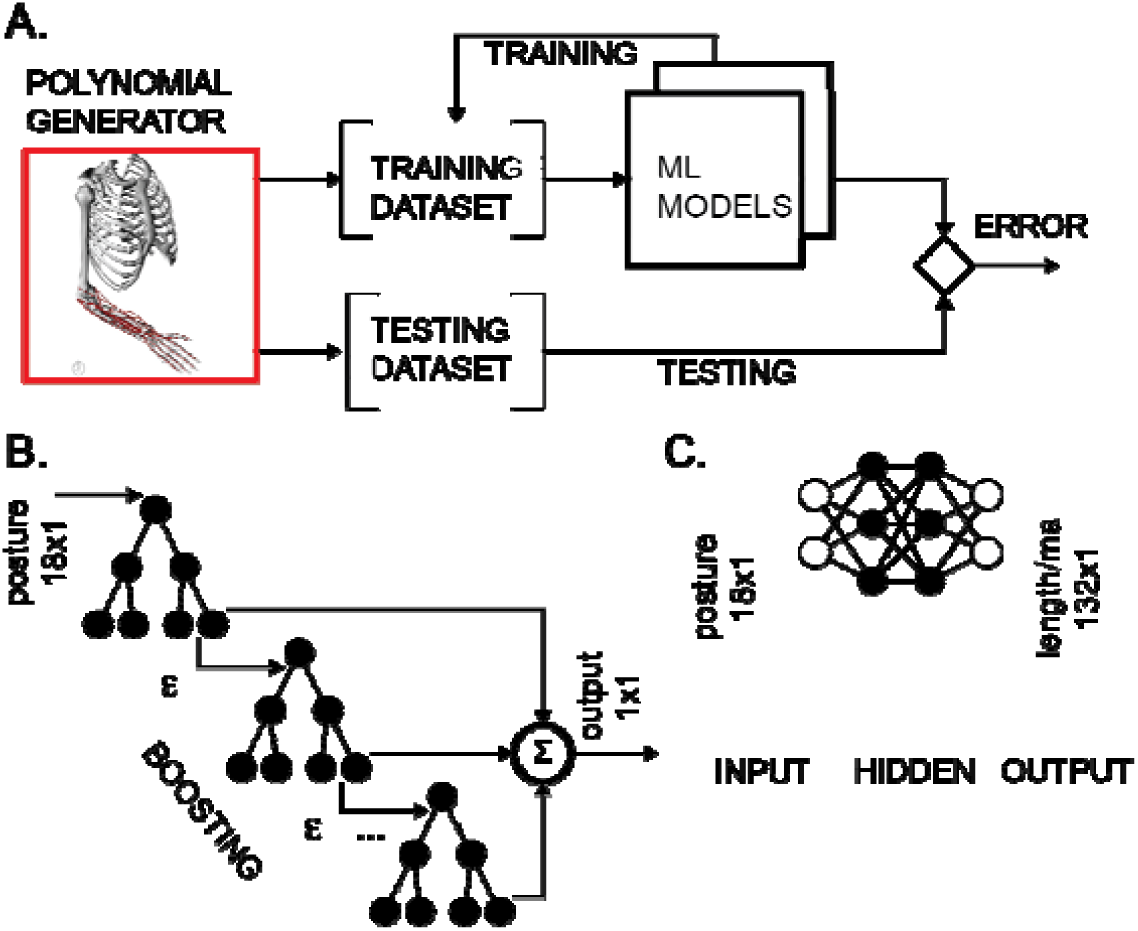
Training and testing of ML models. **A**. The polynomial generator created reference datasets, which were then iteratively used for training and testing. **B**. LGB model with decision binary trees using gradient boosting. The transformation from input postures to output scalar values corresponding to either muscle length or moment arm values was performed in boosting stages to improve accuracy. **C**. ANN with two hidden layers performed transformation for all lengths and moment arms in the model.

Each muscle length and moment arm relationship with posture was fitted with one LGB model. The full arm and hand model was simulated by 33 length and 99 moment arm transformations of 18 dimensional posture input. Three types of hyper-parameters were iteratively optimized prior to training: 1) the number of leaves in a single decision tree (range: 20-100); 2) the minimal number of samples in one leaf (range: 10-100); 3) the maximum tree depth as the number of split levels (range: 1-100). Values for each LGB model were determined iteratively using the Bayesian optimization (Snoek et al., 2012) on training and validation datasets selected as 70% and 30% of a l data, respectively. Other hyper-parameters within LGB models, e.g. the number of weak estimators in boosting (100), were chosen as defaults of Microsoft implementation v.2.2.3 (Ke et al., 2017).

#### Artificial neural network (ANN)

We developed two ANN models to evaluate posture-dependent muscle lengths and moment arms. We selected fully connected feed-forward layers with one input, one output, and two hidden layers (Fig. 2C) (Sazli, 2006) with rectifying linear units as the outputs of every layer. This standard model provides robust gradient propagation with efficient computation (Glorot et al., 2011). Using TensorFlow (Abadi et al., n.d.), we composed our networks consisting of the following number of nodes in input, two hidden, and output layers: [18, 1024, 512, 33] for the approximation of 33 muscle lengths, and [18, 2048, 1024, 99] for the approximation of 99 moment arms.

Xavier initialization method was used to select the initial weights for each layer from the normal distribution with zero mean and its variance as *2/(n*_*in*_*+n*_*out*_), where *n*_*in*_ and *n*_*out*_ were the number of inputs and outputs in this layer (Glorot and Bengio, 2010). The network was trained with the batches of sample data (256 samples) using a gradient based stochastic optimization method minimizing a custom cost function (Kingma and Ba, 2017). We developed the cost function that focused on the performance of the worst approximations evaluated as RMSE of the worst 5% of input-output pairs from each muscle. The scalar cost was evaluated as the mean of all errors within the upper 30% range.

The variable learning rate was used to improve the learning dynamics. The initial rate of 0.001 was reduced by 20% if the measured metric stopped improving after two full training dataset evaluations, or epochs. We have tested additional two manipulations to improve learning. We tested the variation of processing structure to improve the generalization of solution distributed across multiple nodes in the ANN. The model was trained with 50% of the nodes skipped in each evaluation and temporarily and randomly assigned to the dropout layer. In addition, we have also tested the normalization of input samples. However, the improvements due to the additional structure variation and the normalization were marginal, and we chose to exclude these manipulations from the processing pipeline.

The overfitting in training was by comparing the error rates for training (observed) and testing (unobserved) samples. The difference in errors was less than 0.4% for all muscles without the divergence. For example, for as little as 1000 samples, the RMSE of the trained model for *Pronator Teres* length was 78.69% for the training set and 79.02% for the testing set, which indicated the absence of overfitting in the early stage of fitting. The difference between training and testing evaluations remained below 0.1% until the terminal level was achieved.

## Results

Two types of ML models were trained to approximate the musculoskeletal relationships. Our findings detail the training outcome and the training dynamics for learning the transformation from joint posture to muscle lengths and moment arms.

### Estimation of the training dataset size

The selection of the training dataset size for ANN and LGB models is a non-trivial step in the model development. Our source of data was expressed functionally allowing unlimited source of training data. However, the selection of an optimal dataset that captures the relationships without the tendency for overfitting was the initial goal of our development. We used RMSE metric for both length and moment arm models trained with several datasets of incremental size. The relationship between the metric and the dataset size are shown in Figure 3. As the size of the dataset increased logarithmically (from 10^3^ to 10^6^ samples), the training accuracy also increased, with minor improvement in the range above 10^5^ samples. The improvements with the dataset size were not as pronounced showing 1.45% and 1.94% errors with the smallest datasets (10^3^ samples). The improvement curve of LGB is flat, showing no further improvement, after 10^5^ sample size. The performance of the relatively simple (*Biceps Brachii Long Head*) and complex (*Extensor Pollicis Longus*) muscles is illustrated in Fig.3 C and D. These two muscles are on the opposite extremes of complexity expressed as the number of terms required for accurate polynomial fit (Sobinov, et al., 2020). The improvements are qualitatively similar for both of these muscles indicating no strong dependency on muscle path complexity with both types of models. We used the largest dataset size for all further model development described below.

**Figure 3.**
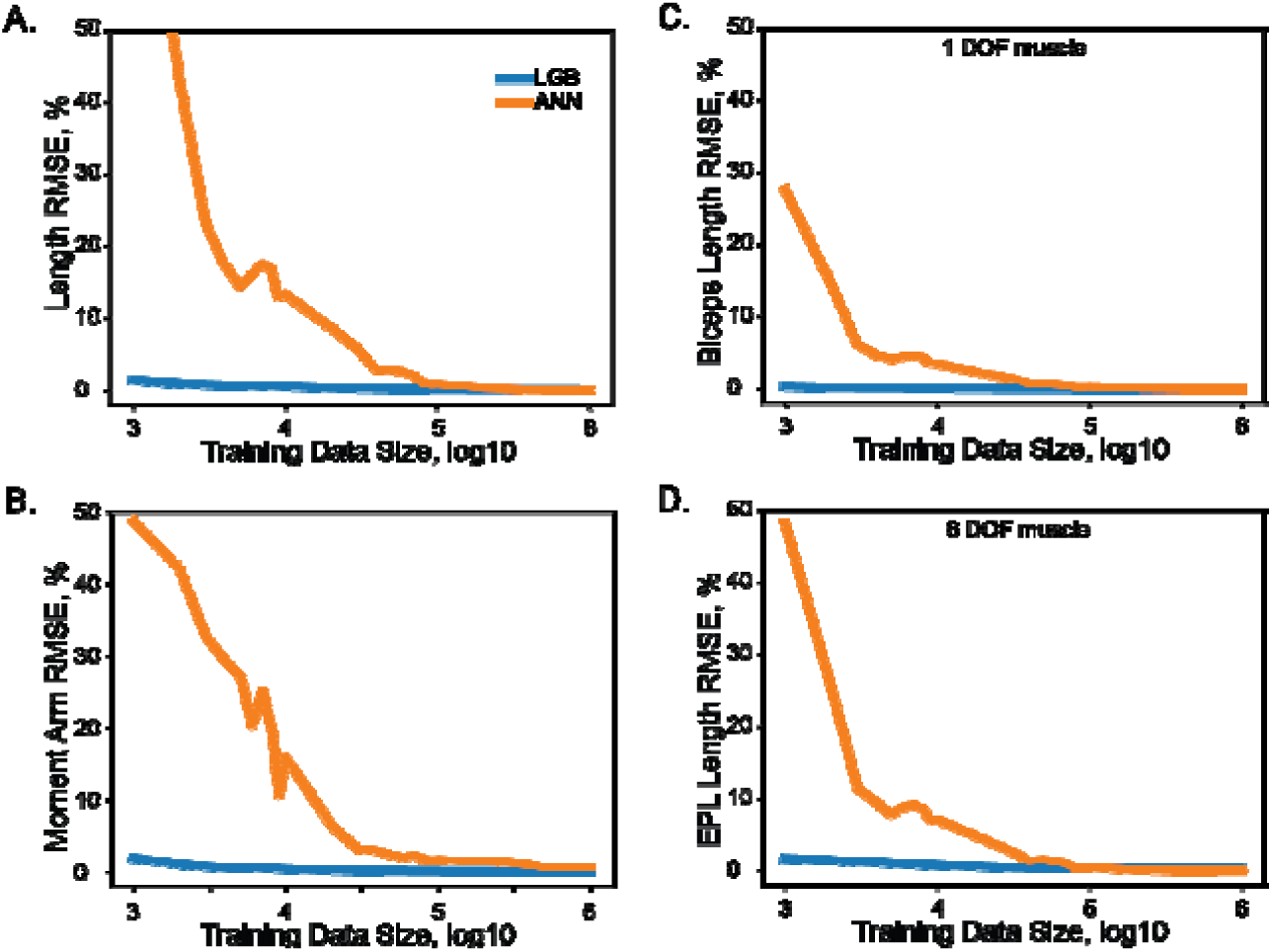
The relationship between model performance (RMSE) and the training dataset size for ANN and LGB models. The length (**A**) and moment arm (**B**) relationships with the size are shown for all muscles. Selected examples of muscle length and the dataset size are shown for *Biceps Brachii Long Head* in **C** and *Extensor Pollicis Longus* in **D**.

### Model accuracy

High accuracy was achieved with both LGB and ANN model types. The distribution of errors is shown in Fig.4 with the histograms of RMSE values for the testing dataset (5*10^4^ samples). The highest achieved performance of ANN models was with 10^7^ with absolute errors at 0.08±0.05% for muscle lengths and 0.53±0.29% for moment arms. Similarly, LGB models generated accurate predictions with large training datasets, 0.18±0.06% and 0.13±0.07% errors, respectively. This is the expected error rate based on our previous analysis (Sobinov et al., 2019).

Overall, the error span did not exceed 0.6% for muscle lengths and 2% for muscle moment arms, shown in Figures 4 and 5. We found no relationship between length and moment arm errors (p=0.746, R^2^=0.003). The level of errors was comparable for both simple and complex muscles (spanning more than 3 DOF), e.g., the errors of ECR_BR (a two DOF muscle) were comparable to those of EDM. The accuracy of LGB and ANN models was comparable. The interquartile ranges (IQR), corresponding to the distance between 25% and 75% level for the distribution of all length error values were 0.075% (ANN) and 0.216% (LGB). The 25-75% IQRs for moment arm errors in Fig. 4B were 0.464% (ANN) and 0.0782% (LGB). Median RMSE values for all models were less than 0.01%. To check if accuracy declines in the extreme postures (1st and 4th quartiles), we repeated testing for errors and found a similar rate of errors, 0.144% and 0.587% for lengths and moment arms.

**Figure 4.**
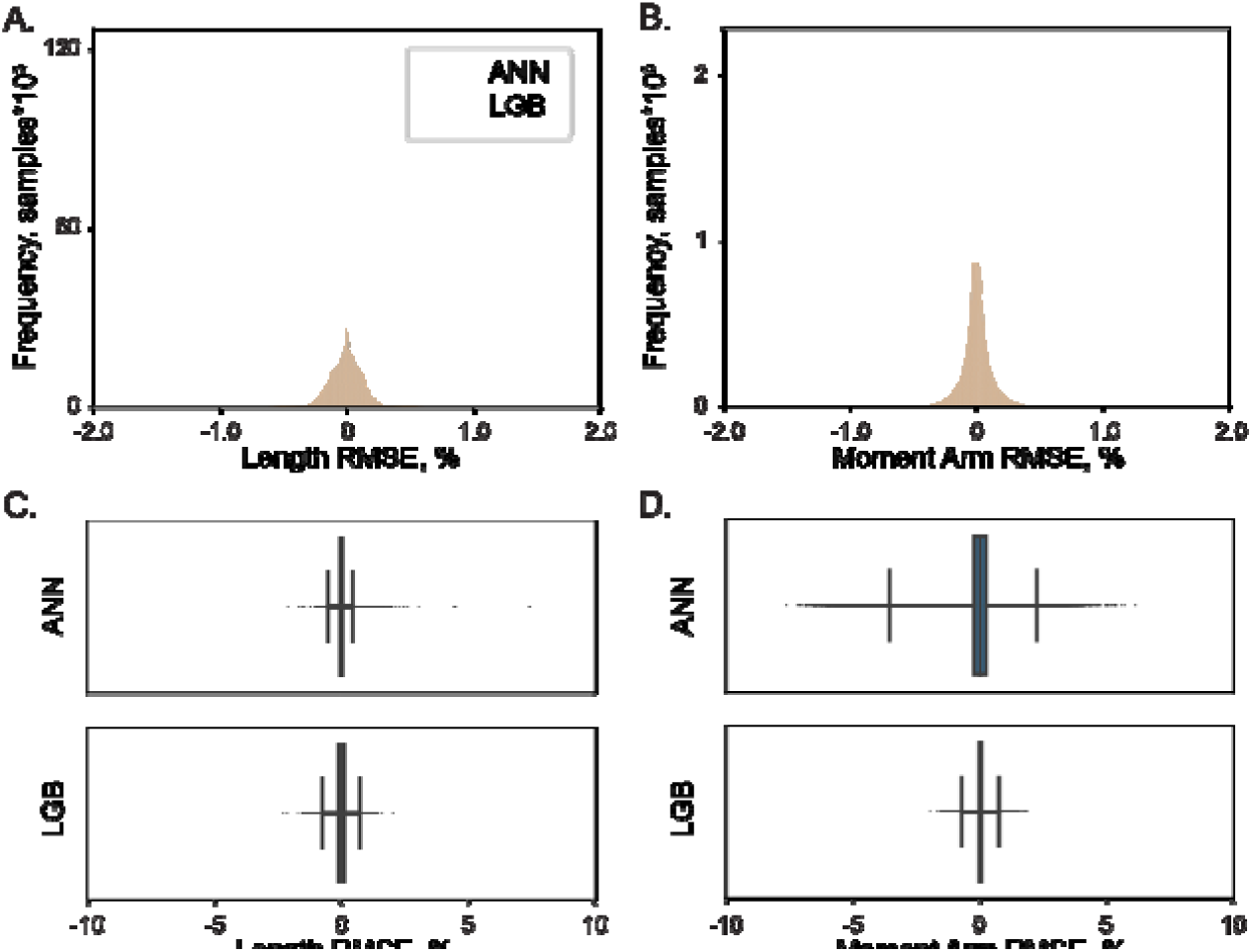
The distribution of normalized errors in the prediction of muscle length (**A**) and moment arms (**B**) are shown for two ML models, LGB (red) and ANN (blue). The distribution of all errors including all the outliers are shown for muscle length (**C**) and moment arms (**D**). The box plots indicate 25% to 75% IQR with whiskers set to cut ±0.1% of the distribution.

**Figure 5.**
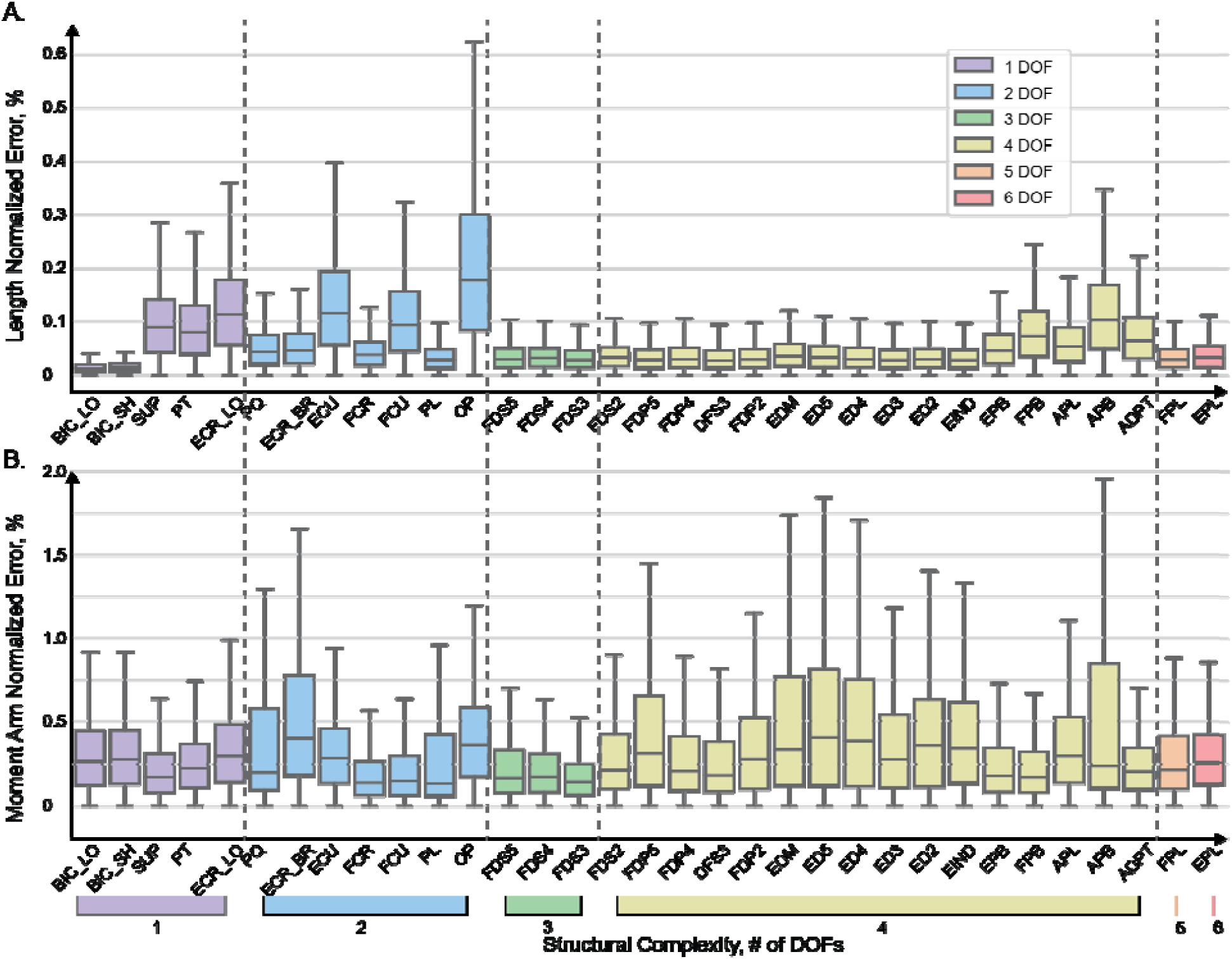
Normalized absolute errors for muscle length (**A**) and moment arms (**B**) evaluated with ANN. The box plots indicate 25% to 75% IQR with whiskers set to cut 1.5% of the distribution.

The distribution of absolute errors is shown for each muscle in Figure 5. The muscles are sorted according to the number of DOFs they actuate with relatively simple muscles on the left and complex (thumb) muscles on the right. Overall, the majority of distributions were below 0.2% for 75% of all vales, as indicated by the top value of the interquartile range in the box plots. The largest length errors were observed in OP, which was also one of the most difficult muscles to design structurally {Boots et al., 2020}. In this muscle the top value (Q3) of the interquartile range was about 0.3%, which corresponds to the error of 0.018 mm. In Fig.5a, the normalized errors of muscle length are presented for each muscle. For all muscles, the most errors (up to 75% of the distributions) are below 0.2%. The prominent exception is OP with the highest normalized errors, which is explained by the minimal full physiological range of only 6 mm. The error of 0.3% in OP length corresponds to the absolute error of 0.018 mm. The errors did not increase with muscle structural complexity. The evaluation errors in muscle length where generally larger in the group of muscles spanning 2 DOF (blue, see *Extensor Carpi Radialis, ERC_LO*) and were comparable to the errors in complex muscles (e.g., *Abductor Pollicis Brevis, APB*).

### Training and evaluation time

The execution times were compared for ANN and LGB models (1.4 GHz Quad-Core 8th-generation Intel Core i5) by measuring the duration of 1000 evaluations (using method *time* from the standard *time* library in Python 3.7). For a given posture, ANN models evaluated both muscle length and moment arms with the combined latency of 1.1±0.6 ms, as compared to 43.1±8.3 ms for LGB models, which were about 39 times slower.

## Discussion

In this study, we solved the musculoskeletal kinematics problem over the full physiological range of limb postures using machine learning approaches. We tested two standard types of models–LGB and ANN–that both accomplished the mapping from limb posture to muscle kinematic state described by multidimensional muscle length and moment arms. LGB and ANN approaches (McGovern et al., 2019; Natekin and Knoll, 2013) were chosen as the machine learning equivalents to the phenomenological model previously developed to approximate posture dependent muscle parameters with polynomial structures (Sobinov et al., 2019). Both machine learning methods produced close approximations with the best results achieved by the ANN approach (RMSE=0.08%) as compared to the LGB approach (RMSE=0.12%) for moment arms. LGB and ANN methods have not been previously demonstrated for the solution of the musculoskeletal kinematics.

### Motor intent decoding

The problem of estimating limb posture from EMG in real-time applications remains to be a current challenge in human-machine interfaces (Kiguchi and Hayashi, 2012). In general, a statistical mapping between posture and recorded activity from descending pathways, nerves, and muscles has been used as the transformation to predict motor intent (Dantas et al., 2019; Lin Wang and Buchanan, 2002; Ting et al., 2019) or to control powered prosthetic limbs or exoskeletal devices (Collinger et al., 2013; Furui et al., 2019; George et al., 2020; Zhang et al., 2017). However, the accuracy of decoding realistic movements remains to be a challenge especially for movements that require dexterous object manipulation (Downey et al., 2017). Many current decoding in brain-computer interfaces assume that neural activity is related to limb end-point velocity and lack description of movement kinetics generated by muscle forces. The resulting movements of prosthetics are typically slower and less robust than natural movements. Up to five (Wendelken et al., 2017) to six (George et al., 2020) arm and hand DOFs can be simultaneously controlled using nerve signals recorded with penetrating electrodes. Accurate but slow movements can be generated for high-dimensional artificial hands with wrist and digits; however, the accuracy is challenged by changing limb posture and orientation. One potential solution is the use of closed-loop control systems that provide not only the forward control of prosthetic, but also incorporate the sensory feedback within neuroprosthetics (Charkhkar et al., 2020; Ganzer et al., 2020; Hughes et al., 2020). The closed-loop control system that takes into account muscle forces will likely require accurate representation of musculoskeletal actions, which are essential for the description of both muscle forces and sensory signals responsible for proprioception.

Another related possible solution that could provide more generalizable control over the nontransparent statistical approaches is the use of mechanistic models that simulate musculoskeletal nonlinearities inherent within the musculoskeletal system. This approach has been demonstrated for the lower-limb (Sartori et al., 2017) and upper-limb (Boots et al., 2017; Crouch and Huang, 2016; Mansouri et al., 2017; Sartori et al., 2018) motor intent decoding and has the major advantage over statistical methods of taking into account kinematic and kinetic relationships between command signals and multisegmented limb posture. Thus, the transformation from the recorded biological signals to the proportional control of limbs should be intuitive, given that the underlying computations are sufficiently accurate and without extensive processing delays.

Together, these possible solutions motivated our exploration and rationale for developing accurate ML methods of approximating the musculoskeletal transformations. We used a model-driven training and testing of machine learning algorithms to approximate posture-dependent changes in muscle lengths and moment arms of distal arm and hand muscles. The previously developed polynomial model provided a functional representation of data across all possible limb postures. Since any volume of data could be generated, we tested the extent of data required to train ANN and LGB models. Figure 3 shows the inverse relationship between the approximation errors and the training dataset size. Surprisingly, the performance of less than 0.5% kinematic errors was achieved with less than 10^6^ samples, which is relatively a low number for the model that includes 33 muscles spanning on average 3 DOFs within a model with 18 DOF. This may indicate that ANN and LGB methods may further generalize the polynomial relationships that required a larger volume of samples to generate functional muscle-posture representations limited to polynomial power terms.

### Computational delays

The transformation of EMG signals into movement rarely accounts for the musculoskeletal anatomy and physiology. This is partly due to the extreme complexity of muscle organization (Gritsenko et al., 2016; Murphy et al., 2018) and nonlinear intrinsic muscle properties that include independent force-length and force-velocity relationships (Zajac, 1989) and less popular in modeling, short-range stiffness, which is a hysteretic force-length property (Cui et al., 2008). The task of simulating muscle force generation requires adequate structural information about muscle paths and posture-dependent changes in moment arms. The development of complex musculoskeletal models has been recently simplified by the dedicated simulation tools for editing and simulating segmental dynamics–OpenSim (Seth et al., 2018), MuJoCo (Todorov et al., 2012), Simscape Multibody (MathWorks, Inc). The challenge remains in collating sufficient datasets of musculoskeletal measurements for creating complex musculoskeletal models and then in testing and validating these models across the full-range of motion to ensure their use in a wide range of applications.

The computational delays of solving the equations of motion governing limb dynamics have previously impeded the application of model-based prosthetic controllers. In particular, the evaluation of muscle force-velocity characteristic may require sub-millisecond latencies to decode accurately rapid movements computed by a physical engine, which requires additional time to execute (∼1ms in Mansouri et al., 2017). Our implementations demonstrated clear speed advantage of ANN over LGB model (about 39x faster) and required about 1 ms on a standard hardware. Further improvements in the performance of ANN may be possible approaching the latencies appropriate not only for the feedforward computations, but also for predictive inverse computations inspired by theoretical neural transformations (Wolpert and Ghahramani, 2000). Another biomimetic feature of ANN model is its potential solution for the necessity to increase computational complexity to accommodate the increase in the size of described structure, typically termed as “the curse of dimensionality”. Sobinov et al. (2019) demonstrated that the typical exponential increase could be replaced with the linear increase in the required number of terms within the polynomial approximations. Here, the same structure of ANN model accommodated accurate calculations for a subset and for the full set of 33 muscles. We hypothesize that the same model can solve models that include other skeletal muscles, if they are relatively similar in anatomical complexity to the muscles represented in our current model.

### Conclusion

We demonstrate in this study the use of two machine learning methods for solving the posture-dependent changes in the musculoskeletal properties essential for limb kinetics. The achieved execution accuracy was adequate with both ANN and LGB models and similar to the original polynomial model. ANN model was 39 times faster than LGB model computing muscle variables in 1.1ms, which is appropriate for real-time control solutions.

